# Notch2 signaling regulates Id4 and cell cycle genes to maintain neural stem cell quiescence in the adult hippocampus

**DOI:** 10.1101/420620

**Authors:** Runrui Zhang, Marcelo Boareto, Anna Engler, Angeliki Louvi, Claudio Giachino, Dagmar Iber, Verdon Taylor

## Abstract

Neural stem cells (NSCs) in the adult hippocampal dentate gyrus (DG) can be quiescent or proliferative, but how they are maintained is largely unknown. With age DG NSCs become increasingly dormant, which impinges on neuron generation. We addressed how NSC activity is controlled and found that Notch2 promotes quiescence by regulating their transition to the activated state. *Notch2*-ablation induces cell cycle genes and markers of active NSCs. Conversely, quiescent NSC-associated genes, including *Id4*, are down regulated after *Notch2* deletion. We found that Notch2 binds the *Id4* promoter and positively regulates transcription. Similar to Notch2, Id4 overexpression promotes DG NSC quiescence and Id4 knockdown rescues proliferation, even when Notch2 signaling is activated. We show that Notch2 regulates age-dependent DG NSC dormancy and Notch2 inhibition rejuvenates neurogenesis in the DG of aged mice. Our data indicate that a Notch2-Id4 axis promotes adult DG NSC quiescence and dormancy.

## Introduction

In the adult mouse brain, neurogenic NSCs persist and are active in the ventricular subventricular zone (V-SVZ) of the lateral ventricles and the subgranular zone (SGZ) of the hippocampal DG (Kempermann, 2011; Zhao et al., 2008). Neurogenesis continues in the brain of human neonates and, although controversial, potentially also in the DG of adults (Boldrini et al., 2018; Sorrells et al., 2018). Most adult murine NSCs in the SVZ and DG niches divide rarely and are in a quiescent state. Once activated, quiescent NSCs enter the cell cycle and start neurogenesis. Activated adult NSCs self-renew to maintain the stem cell pool, generate committed mitotic progenitors to expand the precursor population and differentiate into neuroblasts and glia (Gage, 2000; Ihrie and Alvarez-Buylla, 2011).

Adult neurogenesis in the DG is important for spatial memory but is also implicated in disease including depression and epilepsy (Clelland et al., 2009; Toda et al., 2018; Zhao et al., 2008). To sustain niche homeostasis and continuous neurogenesis, adult NSC maintenance and differentiation are tightly regulated by extrinsic environmental signals and intrinsic genetic factors. Notch signaling is one of the critical niche pathways regulating NSC fate (Ables et al., 2010; Andersen et al., 2014; Basak et al., 2012; Breunig et al., 2007; Ehm et al., 2010; Giachino et al., 2014; Imayoshi et al., 2010; Lugert et al., 2010; Nyfeler et al., 2005). The four mammalian Notch paralogs contain proteolytic cleavage sites essential for maturation and activation upon ligand binding (Mumm and Kopan, 2000). The activated and cleaved Notch intracellular domain (NICD) is released into the cytoplasm, traverses to the nucleus, and interacts with the DNA-binding CSL protein (Rbpj in mice). The Notch/Rbpj transcriptional regulator complex activates expression of target genes including *Hes* and *Hey* family members. The *Hes/Hey* genes encode basic Helix Loop Helix transcription factors that inhibit the expression of the proneural genes and prevent neuronal differentiation while maintaining the NSC pools (Kopan and Ilagan, 2009).

Most analyses of Notch signaling during adult DG neurogenesis investigated the role of Notch1 and its transcriptional effector Rbpj (Ables et al., 2010; Basak et al., 2012; Breunig et al., 2007; Ehm et al., 2010; Imayoshi et al., 2010). Deletion of Rbpj leads to a transient burst in proliferation and production of intermediate progenitors and neuroblasts, and depletes NSCs through accelerated exhaustion of the NSC reservoir (Ehm et al., 2010; Imayoshi et al., 2010; Lugert et al., 2010). Due to its high homology to Notch1, Notch2 was thought to play redundant roles in mediating Notch signaling in the nervous system. However, recent evidence indicates that Notch1 and Notch2 could play different roles, particularly in neurogenesis in the adult V-SVZ. Deletion of Notch1 results in a selective loss of active but not quiescent NSCs, while Notch2 maintains V-SVZ NSC quiescence (Engler et al., 2018). Although Notch2 is highly expressed in the SGZ of the DG, its role in DG neurogenesis is not known.

Here, we investigated the function of Notch2 in adult DG neurogenesis. We found that Notch2 maintains NSC quiescence and its deletion induces proliferation and increases neurogenesis even in the hippocampus of aged mice. Activation of Notch2 signaling blocks transition of DG NSCs from the quiescent to active state but does not drive activated NSCs into quiescence. Notch2 signaling increases *Id4* expression and antagonizes the expression of cell cycle genes to promote NSC quiescence. Notch2ICD interacts with the promoter of the *Id4* gene and promotes transcription. Finally, we show that Id4 expression is sufficient to induce NSC quiescence and is required for Notch2-mediated repression of NSC activation. Our work suggests that a Notch2-Id4 axis controls NSC quiescence and its inhibition could potentially counteract the aging-associated decline in neurogenesis.

## Results

### Notch2 deletion increases progenitor proliferation and neuron generation but results in precocious progenitor exhaustion

In order to investigate the role of Notch2 in adult hippocampal neurogenesis, we initially examined its expression in the SGZ and found it was mainly expressed by GFAP^+^ quiescent NSCs with radial glial morphology (**Figure S1A**). This was confirmed by conditional lineage tracing of *Notch2* expression in Tamoxifen-induced *Notch2::CreER^T2-SAT^*, *Rosa26RCAG::tdTomato* reporter mice (Fre et al., 2011). 1 day after Tamoxifen induction, most tdTomato-labeled cells had a radial glial morphology and expressed GFAP (**Figure S1B**). Quiescent and active DG NSCs express *Hes5* and are labeled in *Hes5::GFP* reporter mice (Lugert et al. 2010). Conversely, NSC activation is associated with expression of Brain Lipid Binding Protein (BLBP) and thus *Hes5::GFP BLBP::mCherry* (*BLBP::mCher*) double reporter mouse separates active and quiescent NSC populations (Giachino et al., 2014). Most quiescent DG NSCs (*Hes5::GFP^+^BLBP::mCher*^-^) expressed both Notch1 and Notch2 whereas only 52.5±1.6% of the active NSCs (*Hes5::GFP^+^BLBP::mCher^+^*) expressed detectable levels of Notch2 protein (**Figure S1C**). In support of these findings, more cells in the GFAP^+^ radial NSC population expressed Notch2 than Notch1 (**Figure S1D**).

To study the function of Notch2 in adult DG NSCs and hippocampal neurogenesis, we conditionally deleted *Notch2* in adult *Hes5::CreER^T2^ Notch2^lox/lox^* mice (*Notch2* CKO), and traced the NSCs and their progeny with a *Rosa26R-CAG::GFP* Cre-reporter (**Figure 1A and S1E**) (Besseyrias et al., 2007; Lugert et al., 2012; Tchorz et al., 2012). 1 day after Tamoxifen induction, *Notch2* mRNA expression by fluorescent activated cell sorted (FACSed) GFP^+^cells from the DG was reduced to approximately 20% of that of control cells. In addition, Notch2 protein was not detectable on GFP^+^ cells of the DG SGZ in *Notch2* CKO, unlike in the control mice (**Figure S1F,G**). 21 days after Tamoxifen treatment, proliferating progenitors (GFP^+^PCNA^+^), newborn neuroblasts (GFP^+^Dcx^+^) and neurons (GFP^+^NeuN^+^) were significantly increased in the *Notch2* CKO (**Figure 1A-D**). Quantification revealed a reduction in the contribution of GFAP^+^ radial glial NSCs (rGFAP) within the GFP^+^ population in the *Notch2* CKO (**Figure 1B-D**). 100 days after Tamoxifen treatment, rGFAP NSCs were significantly lower in *Notch2* CKO than in controls although the number of newborn neurons (GFP^+^NeuN^+^) was still increased (**Figure 1E-G**). The number of newborn neuroblasts (GFP^+^Dcx^+^) and proliferating progenitors (GFP^+^PCNA^+^) were also reduced after 100 days in the *Notch2* CKO (**Figure 1E-G**). Therefore, analysis of the different cell populations in the *Notch2* CKO and controls at day 21 and day 100 after Tamoxifen treatment revealed a rapid reduction in quiescent NSCs and an increase in proliferative progenitors and neuroblasts followed by a putative premature exhaustion of neurogenic NSCs and neurogenic decline within 100 days.

**Figure 1.**
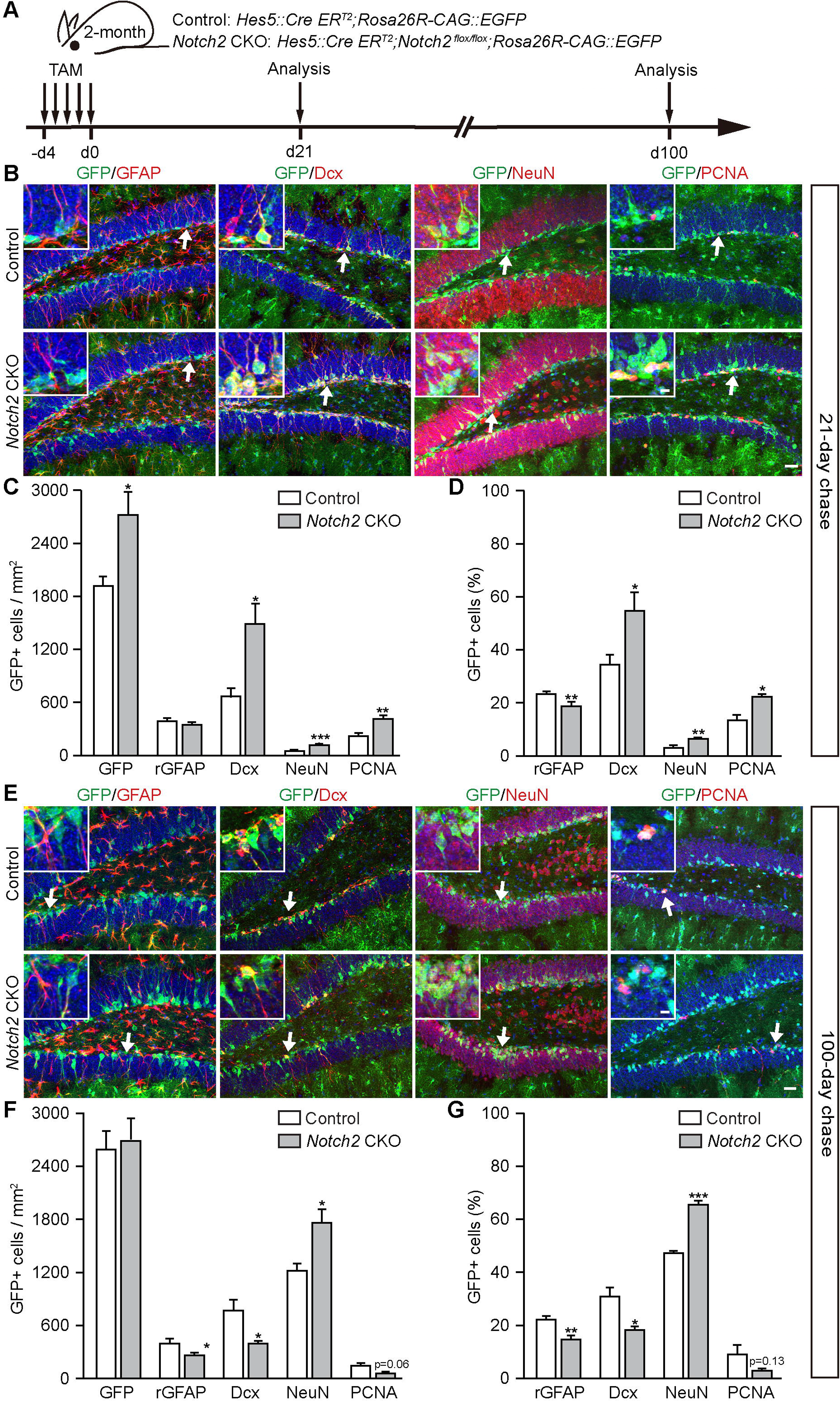
Notch2 maintains NSC quiescence and long-term neurogenesis in the adult hippocampus. **A.** Scheme of the Tamoxifen-induced *Hes5::CreER^T2^*-mediated *Notch2* CKO with 21 day and 100 day lineage chases in adult mice. **B.** Analysis of the progeny of lineage-traced *Hes5::CreER^T2^* expressing control and *Notch2* CKO progenitors (GFP^+^) in the DG 21 days after Tamoxifen treatment. GFAP^+^ - radial quiescent NSCs, Dcx^+^ - newborn neuroblasts, NeuN^+^ - newborn neurons, PCNA^+^ -proliferating progenitors. **C** and **D.** Quantification of GFP, radial GFAP (rGFAP), Dcx, NeuN and PCNA expressing *Notch2* CKO cells (GFP^+)^ in the DG as cells per mm^2^ (**C**) and percentages of genetically labeled GFP^+^ cells (**D**) 21 days after Tamoxifen treatment. **E.** Analysis of the progeny of lineage-traced *Hes5::CreER^T2^* expressing control and *Notch2* CKO progenitors (GFP^+^) in the DG 100 days after Tamoxifen treatment. GFAP^+^ - radial quiescent NSCs, Dcx^+^ - newborn neuroblasts, NeuN^+^ - newborn neurons, PCNA^+^ -proliferating progenitors. **F** and **G.** Quantification of GFP, radial GFAP (rGFAP), Dcx, NeuN and PCNA expressing control and *Notch2* CKO cells (GFP^+^) in the DG as cells per mm^2^ (**F**) and percentages of genetically labeled GFP+ cells (**G**) 100 days after Tamoxifen treatment. Data are means ± SEM (n = 3-4 mice; t-test *p < 0.05, **p < 0.01, ***p < 0.001). Arrows indicate the areas shown in the magnified insets. Scale bar = 30 μm, insets = 6 μm.

### Notch2 activation maintains quiescent NSCs and decreases neuroblast production

We addressed the effects of increasing Notch2 signaling in DG NSCs by conditionally expressing the active form of Notch2 (Notch2ICD) from the *Rosa26* locus (Tchorz et al., 2009). We intercrossed mice to generate *Hes5::CreER^T2^ Rosa26R-CAG::floxed-STOP-Notch2ICD Rosa26R-CAG::GFP* mice and induced Notch2ICD expression in DG NSCs by Tamoxifen treatment. The fate of the *Hes5::CreER^T2^*-derived cells was traced by following GFP expression from the Cre-inducible *Rosa26R-CAG::GFP* reporter (**Figure S2A**). Constitutive activation of Notch2ICD reduced production of GFP^+^ progeny compared to controls (**Figure S2B-D**). Notch2ICD expression also reduced progenitor proliferation (GFP^+^PCNA^+^) and neuroblast (GFP^+^Dcx^+^) production in the DG at 21 days after Tamoxifen treatment (**Figure S2B-D**). The proportion of GFP^+^ *Hes5::CreER^T2^*-derived cells with radial morphology and GFAP expression (rGFAP) was increased in Notch2ICD expressing mice although the total number of labeled rGFAP cells did not change, indicating a reduction in the production of progeny from Notch2ICD expressing NSCs (**Figure S2C,D**). This was also evident from the reduction in the entire lineage including PCNA^+^ progenitors, Dcx^+^ neuroblasts and NeuN^+^ neurons in the DG of Notch2ICD overexpressing mice.

We analyzed early changes in the DG cell lineages 2 days after *Notch2* CKO, or Notch2ICD expression (**Figure S3A**). Notch2ICD expression resulted in a reduction in the number of GFP^+^ cells within 2 days (**Figure S3B-D**). In addition, expression of Notch2ICD reduced proliferation of rGFAP NSCs (GFP^+^PCNA^+^rGFAP^+^) and a concomitant decline in intermediate progenitors (GFP^+^Ascl1^+^), and neuroblasts (GFP^+^Dcx^+^) (**Figure S3B-D**). Conversely, *Notch2* CKO rapidly increased rGFAP NSC proliferation (GFP^+^PCNA^+^rGFAP^+^), intermediate progenitor (GFP^+^Ascl1^+^) production and neuroblast (Dcx^+^) formation (**Figure S3B-D**).

### Notch2 regulates proliferation of quiescent NSCs but not active progenitors

The DG progenitor pool contains quiescent GFAP^+^ radial glial NSCs and actively proliferating GFAP^-^ NSCs and progenitors. To address Notch2 functions in these populations *in vivo*, we infected the DG of adult *Notch2^lox/lox^ Rosa26R-CAG::GFP* mice with *Adeno-gfap::Cre* and Retro-Cre viruses (Rolando et al. 2016). The *Adeno-gfap::Cre* viruses infect and express Cre-recombinase in GFAP^+^ radial glial NSCs of the DG (Merkle et al., 2007). *Adeno-gfap::Cre*-mediated *Notch2* CKO from GFAP^+^ NSCs increased proliferating cells and newborn neurons in the DG compared with control (**Figure 2A,B**). By contrast, Retro-Cre virus infection to delete *Notch2* from mitotically active progenitors in the DG did not affect neuroblast generation (**Figure 2C,D**).

**Figure 2.**
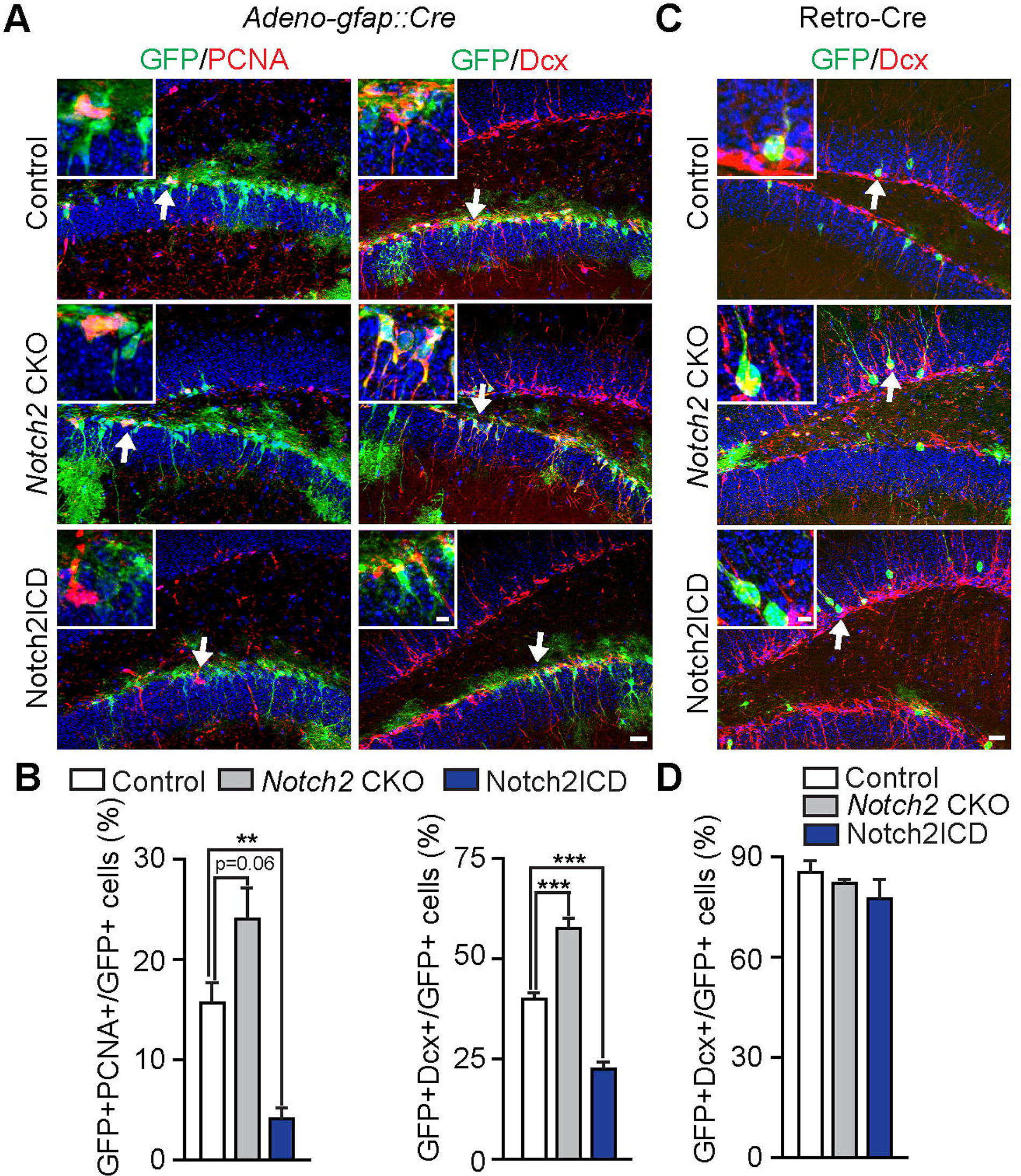
Notch2 signaling regulates quiescent NSCs but not proliferative progenitors. **A.** Analysis of the progeny of lineage-traced *Adeno-gfap::Cre* infected control, *Notch2* CKO and Notch2ICD expressing GFAP^+^ NSCs (GFP^+^) in the DG after 21 days. PCNA^+^ -proliferating progenitors, Dcx^+^ - newborn neuroblasts. **B.** Quantification of PCNA and Dcx expressing control, *Notch2* CKO and Notch2ICD expressing cells (GFP^+^) in the DG 21 days after *Adeno-gfap::Cre* infection. **C.** Analysis of the progeny of lineage-traced Retro*-*Cre infected control, *Notch2* CKO and Notch2ICD expressing mitotic progenitors (GFP^+^) in the DG after 21 days. Dcx^+^ - newborn neuroblasts. **D.** Quantification of Dcx expressing control, *Notch2* CKO and Notch2ICD expressing neuroblasts (GFP^+^) in the DG 21 days after Retro*-*Cre infection. Data are means ± SEM (n = 3-5 mice; t-test *p < 0.05, **p < 0.01, ***p < 0.001). Arrows indicate the areas shown in the magnified insets. Scale bar = 30 μm, insets = 6 μm.

We also infected the DG of *Rosa26R-CAG::floxed-STOP-Notch2ICD Rosa26R-CAG*::GFP with *Adeno-gfap::Cre* and Retro-Cre viruses to address whether activating Notch2 expression affected quiescent NSCs or active mitotic progenitors. *Adeno-gfap::Cre-*mediated activation of Notch2ICD expression in GFAP^+^ DG NSCs reduced the generation of proliferating progenitors and newborn neuroblasts similar to *Hes5::CreER^T2^*-mediated Notch2ICD expression (**Figure 2A,B**). By contrast, Retro-Cre infection and activation of Notch2ICD expression in mitotic progenitors in the DG did not affect proliferation or neuroblast production (data not shown and **Figure 2C,D**). Thus, Notch2 functions to regulate proliferation and neurogenesis in the adult DG are restricted to quiescent NSC and even forced activation in mitotic progenitors does not block neuroblast production.

### Conditional knockout of Notch2 rejuvenates neurogenesis in the aged DG

The DG in aged mice shows a significant decline in neurogenesis, which likely contributes to geriatric cognitive decline (Artegiani and Calegari, 2012; Bizon et al., 2004; Galvan and Jin, 2007; Kempermann et al., 2002). Although proliferating progenitors and newborn neurons are rare in the aged DG, some *Hes5*^+^ dormant NSCs remain (Lugert et al., 2010). We addressed whether Notch2 plays a role in NSC quiescence in the aged mouse brain. We induced *Notch2* CKO in 15-month old *Hes5::CreER^T2^ Notch2^lox/lox^ Rosa26R-CAG::GFP* mice by Tamoxifen treatment and analyzed 21 days later (**Figure 3A**).

**Figure 3.**
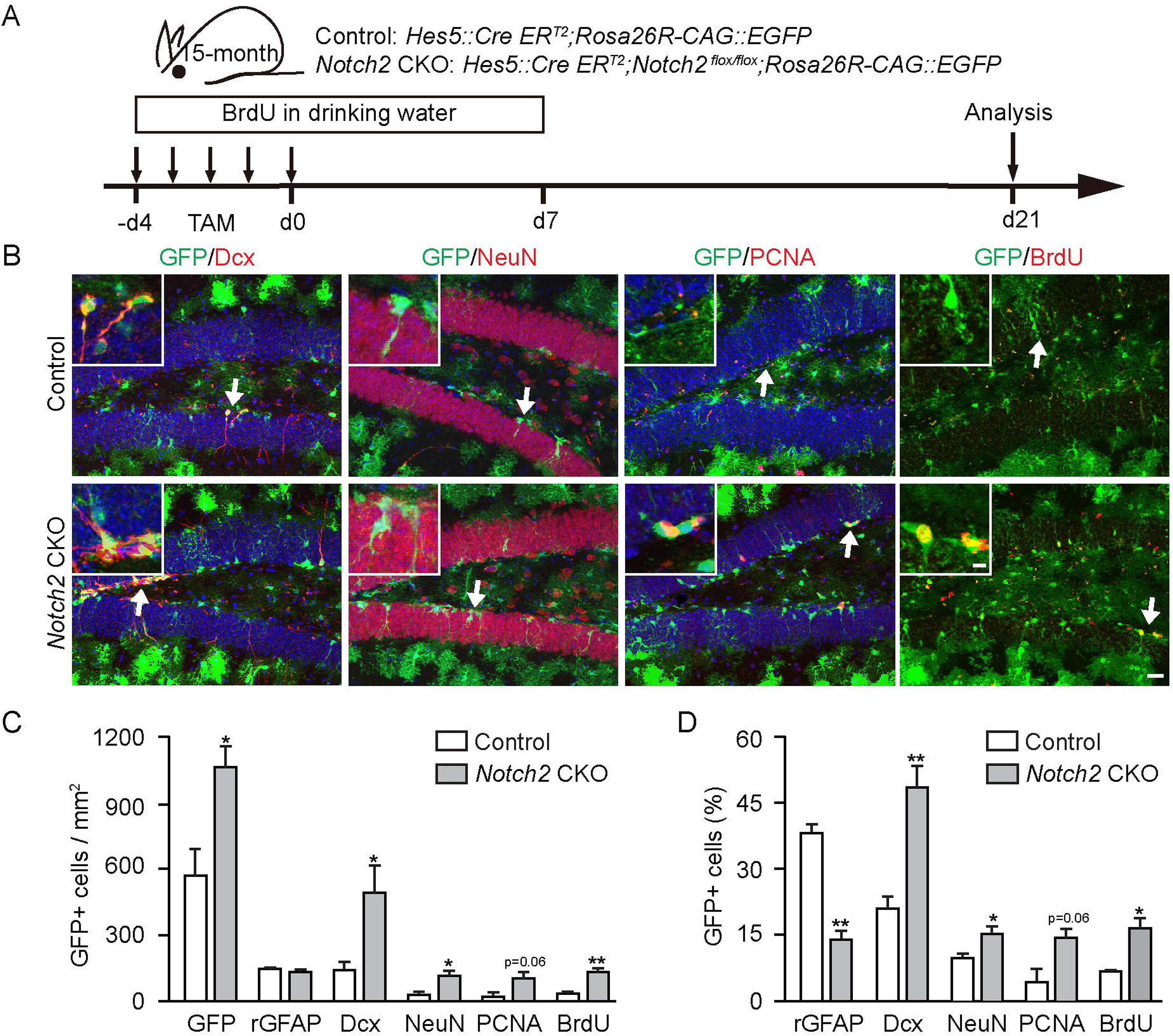
Notch2 deletion rejuvenates neurogenesis in the aged hippocampus. **A.** Scheme of the Tamoxifen-induced *Hes5::CreER^T2^*-mediated control and *Notch2* CKO, BrdU administration and 21 day lineage chase in aged mice. **B.** Analysis of the progeny of lineage-traced *Hes5::CreER^T2^* expressing control and *Notch2* CKO progenitors (GFP^+^) in the DG of aged mice 21 days after Tamoxifen treatment. Dcx^+^ -newborn neuroblasts, NeuN^+^ - newborn neurons, PCNA^+^ - proliferating progenitors, BrdU-labeled cells. **C** and **D.** Quantification of GFP, radial GFAP (rGFAP), Dcx, NeuN, PCNA expressing and BrdU-labeled control and *Notch2* CKO cells (GFP^+^) in the DG of aged mice as cells per mm^2^ (C) and percentages of genetically labeled GFP^+^ cells (**D**) 21 days after Tamoxifen treatment.

*Notch2* CKO in aged mice increased the number of GFP^+^ cells and PCNA^+^ progenitors in the DG indicative of activation of proliferation. To address the proliferation in more detail, we administered BrdU daily to the mice in the first 12 days including during the Tamoxifen treatment and chased 21 days after Tamoxifen treatment. The number of BrdU-labeled newly generated cells increased significantly in the *Notch2* CKO mice compared to controls. In addition, 16.2±2.6% of the GFP^+^ *Hes5::CreER^T2^* -derived cells in the *Notch2* CKO were BrdU-labeled indicating that they were generated through proliferative cell division and not direct differentiation (**Figure 3B-D**). Moreover, *Notch2* CKO from DG NSCs of aged mice induced the production of neuroblasts (Dcx^+^) and neurons (NeuN^+^) compared to control (**Figure 3B,C**). In addition, although the number of rGFAP NSCs did not decrease in the *Notch2* CKO aged mice, they were a reduced population in the *Hes5::CreER^T2^*-derived GFP^+^ cells (**Figure 3C,D**). These results suggest that Notch2 is required for NSC quiescence throughout aging, and blocking Notch2-mediated repression of NSC proliferation in the aged brain rejuvenates neurogenesis.

We addressed the consequences of deleting *Notch2* in adult mice on the aging of the DG. We induced *Notch2* CKO in *Hes5::CreER^T2^ Notch2^lox/lox^ Rosa26R-CAG::GFP* mice by Tamoxifen treatment at 8 weeks of age and then analyzed the mice 10 months later **(Figure S4A)**. Aged *Notch2* CKO animals showed a reduction in rGFAP^+^ NSCs and the proportion of neuroblasts (GFP^+^Dcx^+^) but a higher proportion of neurons (GFP^+^NeuN^+^) at 10 months after *Notch2* deletion (**Figure S4B,C**), thus suggesting that loss of Notch2 had driven quiescent NSCs into neurogenesis and, due to the reduction in NSCs and neuroblasts, neurons (GFP^+^NeuN^+^) were proportionally increased, but the NSC pool was not completely exhausted even after 10 months (**Figure S4A-D**).

### Notch2 down regulates cell cycle genes and active NSC markers, but induces quiescent NSC markers

To investigate the mechanisms by which Notch2 maintains DG NSC quiescence, we performed gene expression profiling of control (*Rosa26R-CAG::GFP*) and *Notch2* CKO GFP^+^ *Hes5::CreER^T2^*-derived cells FACSed from the DG by RNA-seq 1 day after Tamoxifen treatment (**Figure 4A**). We compared three control and *Notch2* CKO samples by supervised variational relevance learning (Suvrel) to identify the largest differences caused by the genotype while reducing technical or individual variations (Boareto et al., 2015). Principle component analysis (PCA) of the RNA-seq data showed clear separation of the control and *Notch2* CKO samples (**Figure 4B**). Among the top 1000 differentially expressed genes between control and *Notch2* CKO, 839 genes were up regulated and 161 genes were down regulated in the *Notch2* CKO samples (**Figure 4C,D** and **Table S8**). The differentially regulated genes were ranked according to Z-score and displayed in a heatmap (**Figure 4D**). GO analysis of the 839 up regulated genes by DAVID (version 6.7) (Huang da et al., 2009) revealed cell cycle and cell division as the most enriched terms and the top 20 regulated genes included cyclins (**Figure 4E,F and Table S8**).

**Figure 4.**
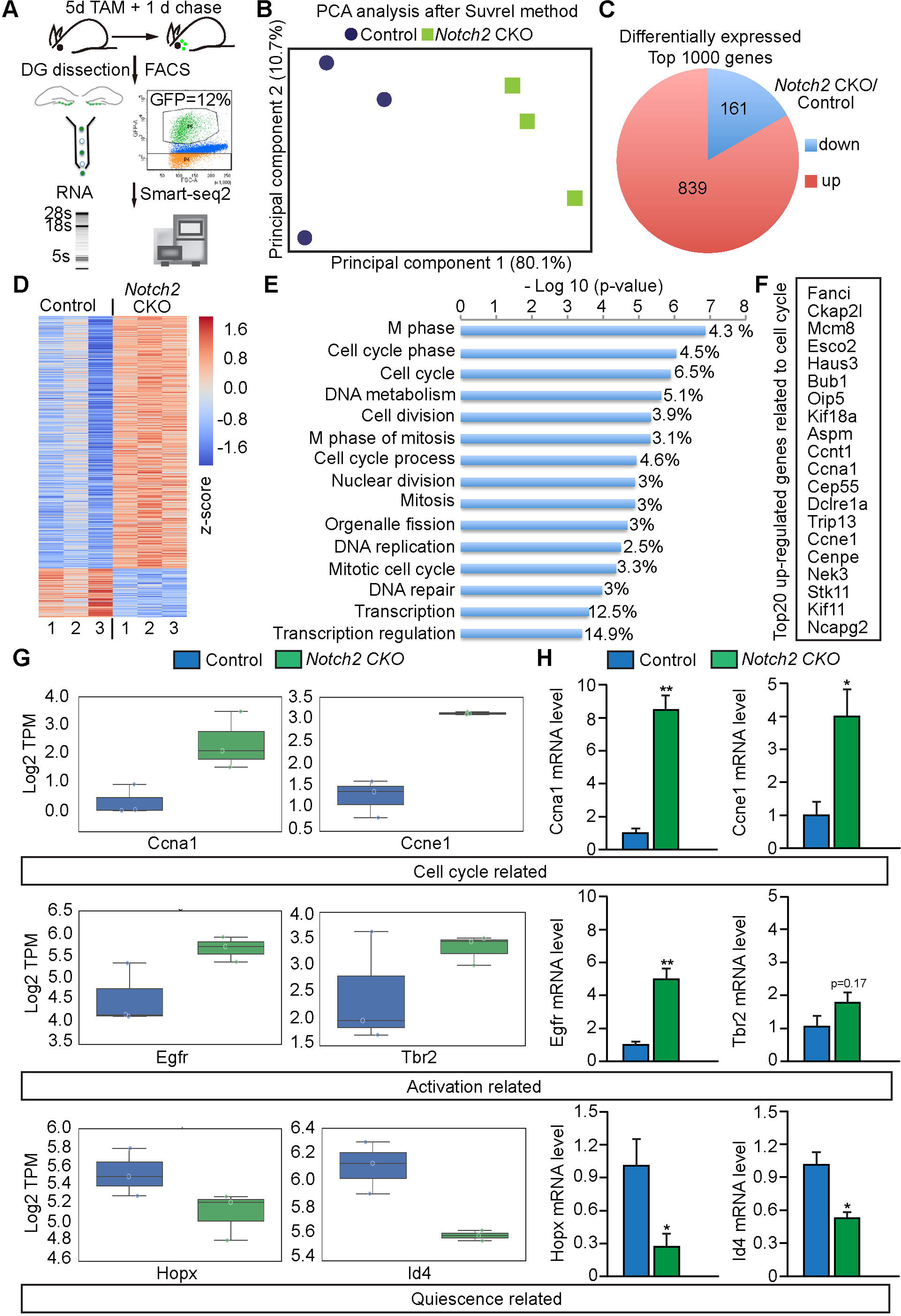
**Notch2 signaling promotes expression of quiescent NSC markers and represses cell cycle-related genes**. **A.** Scheme of sample preparation for RNA-seq analysis. Control (*Hes5::CreER^T2^ Rosa26R-CAG::GFP*) and *Notch2* CKO (*Notch2^flox/flox^*, Hes5::CreER^T2^, *Rosa26R-CAG::GFP*) adult mice were induced with Tamoxifen for 5 days (5d) and sacrificed 24 hours later (1d). GFP^+^ DG cells were sorted by FACS and processed for Smart-seq2 RNA-seq analysis. RNAseq was performed on 3 independent samples of cells pooled from 4-5 mice. **B.** Principle Component Analysis after applying Supervised Variational Relevance Learning (Suvrel) distinguishes control and *Notch2* CKO GFP^+^ *Hes5::CreER^T2^*-derived cell population. **C.** Pie chart of the regulations of the top 1000 differentially expressed genes (*Notch2* CKO/Control). **D.** Heatmap of the top 1000 differentially expressed genes shown as z-score. The ranking of the genes was based on Suvrel (see Methods). **E.** Gene Ontology analysis of the 839 up-regulated genes in *Notch2* CKO cells. **F.** The top 20 differentially expressed cell cycle-related genes from the GO analysis. **G.** Expression of the cell cycle-related genes Ccna1 and Ccne1, the epidermal growth factor receptor (EGFR), the transcription factors Tbr2 and Hopx and Id4 based on counts from the RNA-seq expression analysis. **H.** Quantitative PCR analysis of gene expression on independent samples of FACSed GFP^+^ DG cells from control and *Notch2* CKO mice. Cell cycle-related genes (*Ccna1* and *Ccne1*), NSC activation regulators (*Egfr* and *Tbr2*) and NSC quiescence-related genes (*Hopx* and *Id4*). Expression levels in (**G)** are presented as the log2(TPM+1). Data in (**H**) are presented as mean ± SEM (n = 3 biological replicates; t-test *p < 0.05, **p < 0.01).

We focused on the differentially regulated genes in our RNA-seq dataset that have been previously linked to NSC quiescence and activation (Shin et al., 2015). After *Notch2* deletion, quiescent NSC-associated genes, including *Hopx* and *Id4* were down regulated (**Figure 4G**).By contrast, genes associated with NSC activation or proliferative progenitors including *Egfr* and *Tbr2* were up regulated (**Figure 4G**). We confirmed the regulation of these genes by quantitative RT-PCR analysis on independent RNA samples (**Figure 4H**). Together, these data suggest that loss of Notch2 drives NSCs into an active and proliferative state and genes associated with mitosis are up regulated whereas genes associated with quiescent NSCs are down regulated.

### Notch2 directly regulates the *Id4* gene

To address whether some of the genes regulated after *Notch2* CKO are potential direct targets of Notch2 signaling, we performed Notch2ICD chromatin immunoprecipitation (ChIP) assay from DG NSCs (**Figure 5A**). We isolated and cultured adult DG NSCs from control (*Rosa26R-CAG::GFP*) and conditional Notch2ICD expressing mice (*Rosa26RCAG::floxed-STOP-Notch2ICD Rosa26R-CAG::GFP*) (**Figure 5A**) (Rolando et al. 2016). The DG NSCs were infected *in vitro* with *Adeno-Cre* viruses to induce gene recombination resulting in GFP expression from the *Rosa26R-CAG::GFP* locus and myc-tagged Notch2ICD expression from the *Rosa26R-CAG::floxed-STOP-Notch2ICD* locus (**Figure 5A**).

**Figure 5.**
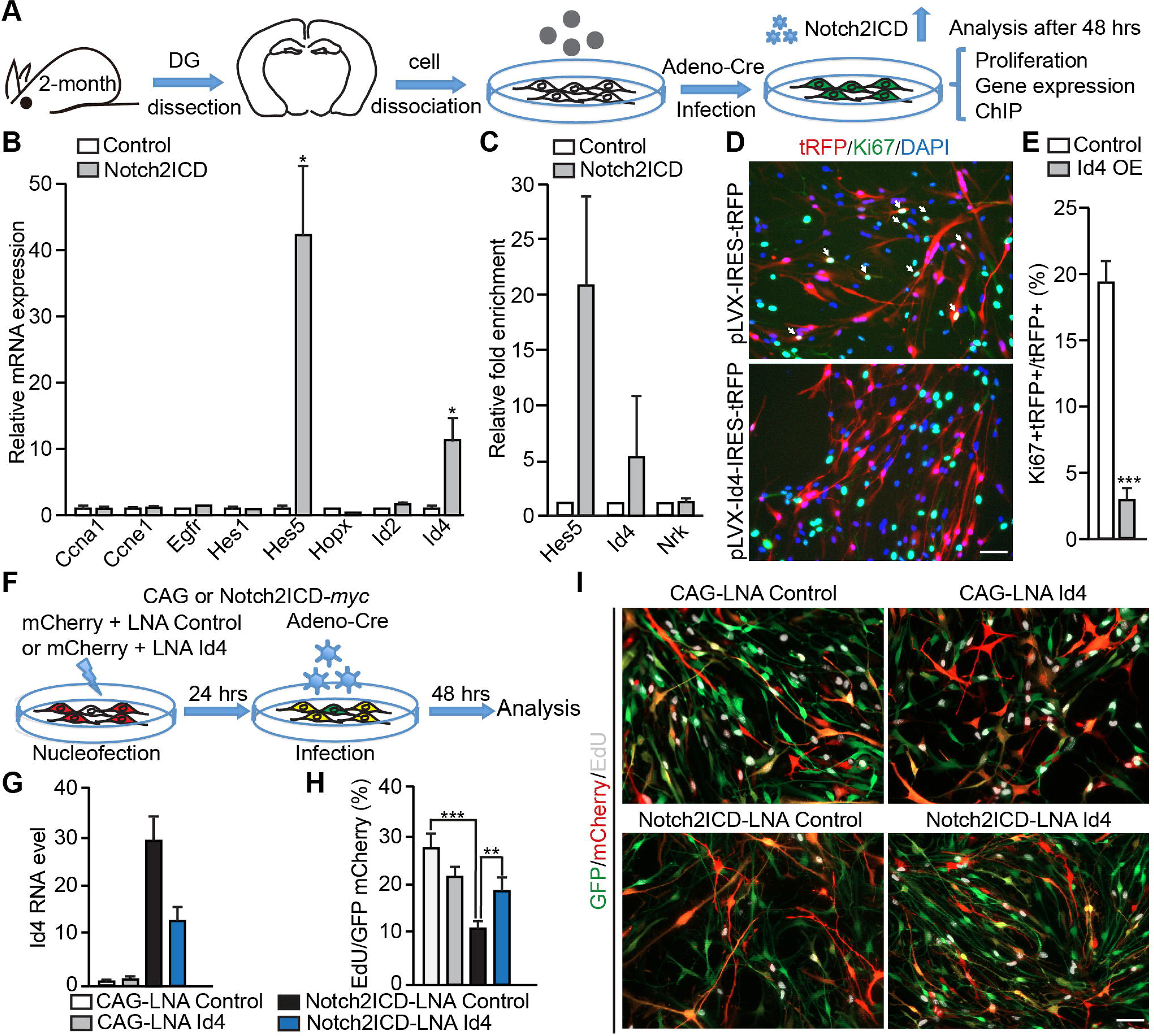
Notch2 directly regulates Id4 to promote NSC quiescence *in vitro*. **A.** Scheme of *in vitro* analysis of Notch2ICD-overexpressing DG NSCs. Adherent DG NSC were derived from adult mice carrying *Rosa26R-CAG::floxed-STOP-Notch2ICD* and *Rosa26R-CAG::GFP* allele alone (control). The cells were infected with *Adeno-Cre* viruses and recombination controlled by the expression of GFP. NSC cultures were collected and used for phenotypic, gene expression and ChIP-PCR analysis. **B.** Relative mRNA expression of *Ccna1, Ccne1, Egfr, Hes1, Hes5, Hopx, Id2 and Id4* in control and Notch2ICD expressing DG NSCs 48 hours of *Adeno-Cre* infection. *Hes* genes and other putative Notch2-regulated targets selected from *Notch2* CKO RNA-seq data were examined. **C.** Chromatin immunoprecipitation of *Rosa26R-CAG::floxed-STOP-Notch2ICD*, which carries a Myc epitope with anti-Myc antibodies. Notch2ICD binds the promoters of *Hes5* and *Id4*. Expression is relative to Myc antibody ChIP from control cells. *Nrk* promoter is not bound by Notch2ICD and was used as a negative control. The error bar represents SEM from three independent ChIP experiments with independent samples. **D.** Overexpression of Id4 IRES tRFP in adult DG NSC (tRFP: red) or tRFP alone (IRES tRFP) and immunostained for the proliferation marker Ki67. **E.** Quantification of Ki67^+^ cells by Id4 IRES tRFP overexpressing (Id4 OE) and IRES tRFP only expressing control cells. **F.** Scheme of the analysis of Id4 function in Notch2ICD-induced NSC quiescence *in vitro*. Id4 KD with locked nucleic acid (LNA) probe and Notch2ICD expression 24 hours later by *Adeno-Cre* infection of DG NSCs from *Rosa26R-CAG::floxed-STOP-Notch2ICD Rosa26RCAG::GFP* (*Notch2*ICD) and *Rosa26R-CAG::GFP* (control) mice. NSCs were analyzed 48 hours after *Adeno-Cre* infection. LNA transfection was monitored by expression of *CMV*::mCherry (mCherry). **G.** Quantitative PCR analysis and relative expression of Id4 expression from sorted GFP mCherry double positive cells. **H.** Quantification of EdU incorporation by Notch2ICD expressing (GFP^+^) LNA transfected (mCherry) cells. **I.** EdU incorporation by Notch2ICD expressing (GFP^+^) and control (GFP^-^), Id4 KD cells (mCherry^+^).

Control and Notch2ICD expressing DG NSCs expressed the NSC/NPC markers Sox2 and Nestin (**Figure S5A,B**). We addressed proliferation of the control and Notch2ICD expressing DG NSCs by BrdU incorporation assay and Ki67 immunostaining (**Figure S5C,D**). Notch2ICD expressing DG NSCs showed a reduction in BrdU incorporation after a 4-hour pulse prior to fixation and fewer cells expressed Ki67 suggesting Notch2ICD expressing cells preferentially exited cell cycle compared to controls (**Figure S5C,D**).

We examined the expression level of several candidate genes selected from RNA-seq data in Notch2ICD overexpressing NSC cultures. Not surprisingly, *Hes5*, a canonical Notch target gene, was up regulated by Notch2ICD expressing NSCs (**Figure 5B**). However, *Id4* mRNA expression was also dramatically up regulated (**Figure 5B**). This finding was in agreement with our finding that *Notch2* CKO leads to *Id4* mRNA down regulation (**Figure 4G,H**). The other cell cycle genes and stem cell markers analyzed were not significantly changed (**Figure 5B**). Thus, our data indicate that Notch2 strongly regulates *Id4* expression at the transcriptional level.

We addressed whether the regulation of *Id4* expression by Notch2 is potentially direct. We identified two Rbpj-binding motifs between base pairs 496 and 699 upstream of the *Id4* transcription start site (**Figure S5E,F**). Therefore, we analyzed whether Notch2ICD could directly bind this region of the *Id4* promoter. Using the *Hes5* gene as a positive control and an intergenic region of the *Nrk* gene as a negative control, we performed Notch2ICD-myc ChIP-qPCR from the DG NSCs to test for enrichment of these sites. The *Id4* promoter flanking region was enriched approximately 5-fold in Notch2ICD expressing cells compared to control cells or to the intergenic *Nrk* region (**Figure 5C**). The *Hes5* promoter was also specifically enriched in the Notch2ICD ChIP from DG NSCs supporting activation of canonical Notch signaling upon expression of Notch2ICD (**Figure 5C**). These data indicate that the *Id4* gene is a direct target of Notch2 signaling.

### Id4 promotes DG NSC quiescence

To test if Id4 regulates NSC quiescence, we transfected adult DG NSCs *in vitro* with pLVX-Id4-IRES-TagRFP expression vectors to overexpress Id4 and traced the cells 48 hours later with the TagRFP. Id4 overexpression resulted in a significant reduction in the expression of Ki67 by NSC and thus increased cell cycle exit (**Figure 5D,E**). This suggested that Id4 expression can drive NSCs into a quiescent state, and this was consistent with Notch2 signaling inducing *Id4* expression in quiescence DG NSCs. We asked whether Id4 plays a role in Notch2-induced DG NSC quiescence. Therefore, we knocked down *Id4* expression in *Rosa26R-CAG::floxed-STOP-Notch2ICD Rosa26R-CAG::GFP* DG NSCs with locked nucleic acid (LNA) probes and induced Notch2ICD expression by *Adeno-Cre* infection 24 hours later (**Figure 5F**). 48 hours after *Adeno-Cre* infection, *Id4* mRNA expression was reduced **(Figure 5G)** and EdU incorporation increased in *Id4* LNA-treated Notch2ICD overexpressing cells (**Figure 5H,I**). Hence, *Id4* knockdown resulted in a partial rescue of the proliferative defect in DG NSCs induced by Notch2ICD expression.

### Notch2 regulates Id4 expression in DG NSCs

To further validate the regulation of the Notch2-Id4 cascade *in vivo*, we co-stained for Id4 and GFAP in control, *Notch2* CKO and Notch2ICD expressing adult DGs 2 and 21 days after Tamoxifen administration (**Figure 6A-E**). Id4 is expressed by radial GFAP^+^ NSCs in the DG SGZ. GFAP^+^Id4^+^ radial NSCs were significant reduced after *Notch2* CKO at both 2 and 21 days post-Tamoxifen treatment indicating maintenance of Id4 by Notch2 signaling (**Figure 6B**). In support, the Id4^-^ NSC population (GFAP^+^Id4^-^) was dramatically increased 2 days after *Notch2* CKO, but decreased at 21 days suggesting precocious exhaustion of the population (**Figure 6C**). Conversely, the GFAP^-^Id4^-^ differentiated cell populations were not dramatically changed at 2 days post-*Notch2* CKO, but were significantly increased by 21 days (**Figure 6D**). Thus, the loss of Id4 correlates with a transient enrichment in GFAP^+^Id4^-^ cells at 2 days and accumulation of GFAP^-^Id4^-^ populations at 21 days in the DG of *Notch2* CKO. The proliferating radial NSC population (GFAP^+^PCNA^+^) paralleled the transient increase in Id4-GFAP^+^ NSCs (**Figure S3C,D**). Following *Notch2* CKO, the proportion of GFP^+^GFAP^+^ *Hes5::CreER^T2^*-derived NSCs that expressed Id4 was significantly reduced at 2 days (**Figure 6E**), lending further support to the hypothesis that *Notch2* CKO leads to loss of Id4 from radial NSCs. At 21 days post-*Notch2* CKO the change in GFAP^+^Id4^+^ radial NSCs was no longer significant, mainly due to the reduction in the NSC population (**Figure 6E** and **Figure 1D**). In addition, Notch2 activation by expression of Notch2ICD prevents the transition from radial GFAP^+^Id4^+^ NSCs to GFAP^-^Id4^-^ progeny, thus maintaining the NSC pool (**Figure 6E**). From these data, we conclude that Notch2 directly regulates Id4 to preserve the NSC pool *in vivo* (**Figure 6F**).

**Figure 6.**
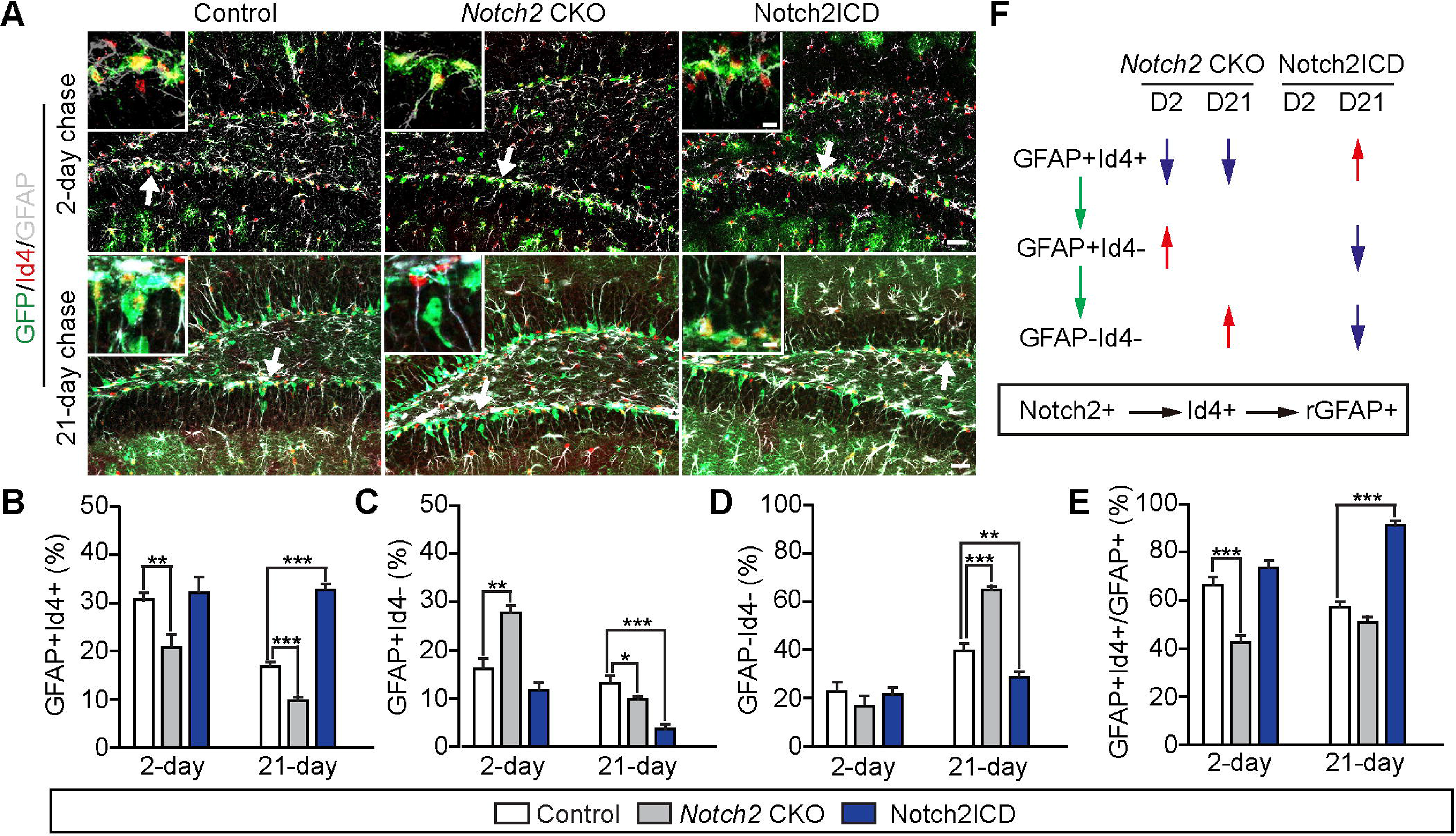
**Notch2 directly regulates Id4 to promote NSC quiescence *in vivo***. **A.** Expression of Id4 and GFAP by progeny of the *Hes5::CreER^T2^* NSCs in the DG of *Hes5::CreER^T2^ Rosa26R-CAG::GFP* control, *Notch2* CKO and Notch2ICD expressing adult mice at 2 days and 21 days after 5 day Tamoxifen treatment. **B-E.** Quantification of GFAP, Id4 expressing radial NSCs in the DG of control, *Notch2* CKO and Notch2ICD overexpressing mice 2 days and 21 days after Tamoxifen treatment. **F.** Summary of the changes in GFAP, Id4 expressing cells in the DG of control, *Notch2* CKO and Notch2ICD overexpressing mice 2 days (D2) and 21 days (D21) after Tamoxifen treatment. Data are means ± SEM (n = 9-11 sections from 3-4 mice; t-test *p < 0.05, **p < 0.01, ***p < 0.001). Arrows indicate the areas shown in the magnified insets. Scale bars = 30 µm, insets = 6 μm.

## Discussion

Here we demonstrated that Notch2 regulates NSC quiescence in the adult DG. Loss of Notch2 signaling rapidly triggers cell cycle entry of radial NSCs and induces neurogenesis. In addition, we show that Notch2 regulation of DG NSC quiescence is responsible for NSC dormancy in the aged mouse brain (Figure 7). Therefore, the transition of DG NSCs to a dormant state in geriatric mice uses the same Notch2 signaling pathway used to maintain a large proportion of the adult DG NSCs. These findings in the DG are consistent with our recent demonstration of Notch2 function in the adult V-SVZ (Engler et al. 2018). In the DG, as in the V-SVZ, Notch2 is primarily responsible for preventing NSCs from entering the cell cycle. Conversely, Notch1 regulates maintenance of the activated NSC pool presumably by promoting self-renewal and preventing differentiation of mitotic NSCs (Basak et al. 2012, Bruenig et al. 2007). Our data suggest aging in the DG NSC pool reflects an imbalance between Notch signaling and proneurogenic cues. An age-related imbalance in Notch and niche signals has also been proposed to regulate muscle satellite cell activation and maintenance (Carlson et al., 2009; Conboy et al., 2003). However, in skeletal muscle, activation of Notch signaling is critical to rejuvenate satellite cells (Carlson et al., 2009; Conboy et al., 2003).

**Figure 7.**
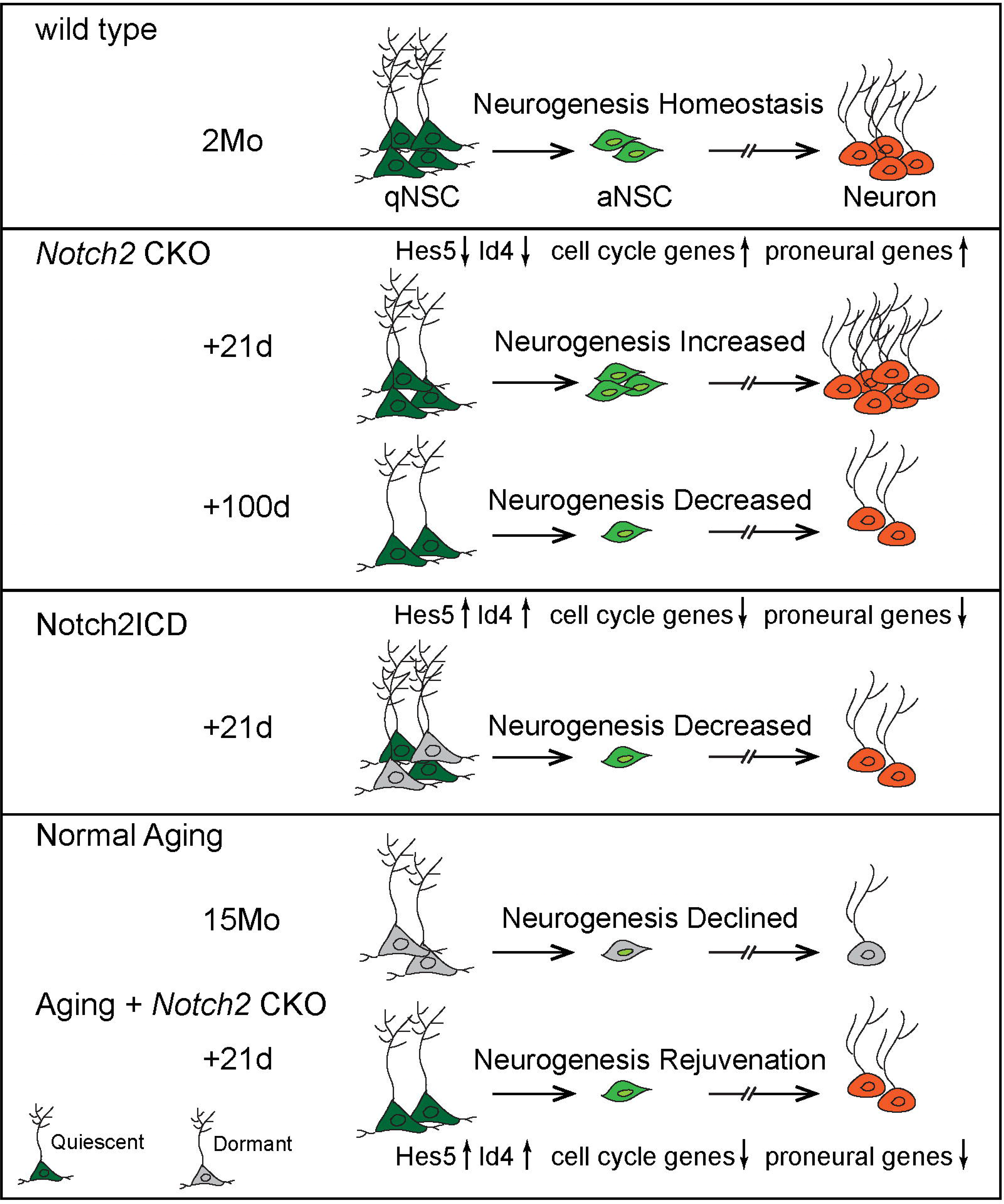
**The model showing Notch2 maintenance of NSC quiescence in the adult hippocampus through regulating Id4 and cell cycle genes**. In the adult hippocampus, deletion of Notch2 down-regulates *Hes* and *Id* genes, and induces cell cycle and proneural genes. This leads to a rapid increase in proliferation and production of newborn neurons (day 21), but exhaustion of the stem cell pools and impaired long-term neurogenesis (day 100). Conversely, activation of Notch2 signaling up regulates *Hes* genes and *Id* genes, and down regulates cell cycle and proneural genes. These maintain NSCs in a quiescent state and reduce neurogenesis. In the aged hippocampus, deletion of Notch2 can rejuvenate neurogenesis by activating dormant NSCs.

In the adult DG, Notch2 regulates NSC activity partially through the activation of Hes protein expression to repress the proneural transcription factor Ascl1, which is required by DG NSCs for activation (Andersen et al. 2014). Oscillatory expression of Ascl1 promotes NSC entry into cell cycle and maintained high-level expression drives differentiation. However, Ascl1 is also controlled at the post-translational level by the E3 ubiquitin ligase Huwe1, which promotes Ascl1 degradation and prevents NSC differentiation and transition from an active to quiescent state (Urban et al. 2016). Interestingly, Huwe1 does not regulate the transition of quiescent NSC to activity. The control of NSC maintenance by Ascl1 seems to be critical and hence Ascl1 is regulated at multiple levels.

We further show that the inhibitor of DNA binding protein Id4 has an important effect to mediate Notch2 regulation in the maintenance of DG NSC quiescence. Notch2 regulates DG NSC cell cycle entry and Id4 is a key mediator of this effect. Notch2 binds the promoter of the *Id4* gene to activate expression. Similarly, Notch1ICD has been shown to bind the *Id4* promoter in embryonic neural progenitors (Li et al., 2012). Interestingly, in the developing nervous system of *Xenopus laevis*, Notch signaling represses *Id4* expression and in mice Id4 regulates neural progenitor proliferation and differentiation (Bedford et al., 2005; Liu and Harland, 2003; Yun et al., 2004). This may reflect the different mitotic states and proneural factors used during adult and embryonic neurogenesis. Id4 expression in cultured DG NSCs reduces cell cycle and knockdown of Id4 is sufficient to reverse the cell cycle effects of *Notch2* CKO on DG NSCs. Thus, we suggest that Id4 is a downstream target of Notch2 signaling, and Id4 is indispensable to prevent NSC activation and differentiation. Ids can release Hes auto-repression thereby stabilizing high level Hes protein expression that in turn represses proneural factor genes (Bai et al., 2007). Id proteins are also known to form inactive dimers with E-box proteins and thereby sequester these coactivator partners of the proneural factors, including Ascl1. In the absence of E-box protein, Ascl1 is unable to activate target gene transcription (Bertrand et al., 2002; Jen et al., 1992). Ascl1 induces differentiation in the embryonic nervous system and cell cycle entry of quiescent adult NSCs (Andersen et al., 2014). Moreover, recent data show that as a result of binding and sequestering the E-box protein E47, Id4 enhances the degradation of Ascl1, thereby maintaining DG NSC quiescence (Blomfield et al., 2018). These observations all support our findings that Notch2 conveys quiescence to NSCs through Id4. This stem cell maintenance activity of Id4 in the DG is somewhat in contrast to its role during embryonic brain development where it promotes NSC proliferation (Bedford et al., 2005; Yun et al., 2004).

Id proteins are downstream effectors of the TGF-βBMP pathway (Hollnagel et al., 1999). BMP signaling has been show to repress DG NSC proliferation *in vitro* and thus potentially regulates DG neurogenesis *in vivo* (Martynoga et al., 2013; Mira et al., 2010). It is currently unclear whether BMP signaling is sufficient for DG NSC quiescence in the adult brain. Thus, our finding that Notch2 regulates Id4 expression in NSCs in the adult DG indicates a potential synergy between Notch2 and BMP. Partial redundancy of Notch2 with other pathways is likely as some radial NSCs remained in the *Notch2* CKO DG. Either the hippocampus contains different NSC populations with differing dependence on Notch2, or other pathways can partially compensate for the loss of Notch2, and this could include BMPs.

Recent single cell RNA-seq data also suggest that both Id4 and Notch2 are highly expressed by adult hippocampal quiescent NSCs (Llorens-Bobadilla et al., 2015; Shin et al., 2015). Therefore, our demonstration of a direct link between Notch2 and Id4 expression in NSCs combines two key pathways associated with NSC regulation in the adult brain. In support, we recently generated a mathematical model based on RNA-seq data from NSCs and their progeny, which predicted interactions between Notch and Id factors as a foundation for NSC maintenance and neurogenesis (Boareto et al., 2017). Taking into consideration that maintenance of adult NSCs requires orchestration of complex regulatory networks, our work emphasizes Notch signaling, Id genes, and cell cycle regulators as critical components in these NSC regulatory networks. Since quiescent and active NSCs have unique gene signatures and landscapes, it remains to be shown how Notch2 and Id4 control the expression of other stem cell markers and proneural genes to maintain the NSC pool (Llorens-Bobadilla et al., 2015; Shin et al., 2015).

## Author Contributions

Conceptualization, R.Z., and V.T.; Investigation, R.Z., M.B., A.L., and A.E.; Methodology and scientific advice, C.G., and D.I.; Writing – Original Draft, R.Z. and V.T.; Writing – Review and Editing R.Z., M.B., A.E., A.L., C.G. D.I. and V.T.; Funding Acquisition D.I., R.Z. and V.T.; Project Administration V.T.

## Acknowledgments

We would like to thank Dr. Jan S. Tchorz for providing *Rosa26R-CAG::floxed-STOP-Notch2ICD* mice, Dr. Spyros Artavanis-Tsakonas for providing *Notch2::CreER^T2-SAT^ Rosa26R-CAG::tdTomato* mouse, Drs. Kirsten Obernier and Arturo Alvarez-Buylla for providing the *Adeno-gfap::Cre* virus, Dr. H. R. MacDonald for providing the Notch1 and Notch2 antibodies and Fred Sablitzky for providing the Id4 cDNA clone. We thank the members of the Taylor lab for critical reading and correction of the manuscript, and Frank Sager for excellent technical assistance. We thank Dr. Christian Beisel of Genomics Facility Basel of D-BSSE for NGS RNA sequencing, the animal core facility of the University of Basel and BioOptics Facility of the Department of Biomedicine for support. We thank Francois Guillemot, Noelia Urban and Isabelle Blomfield for sharing unpublished data. This work was supported by the SystemsX.ch project NeuroStemX (51RT-0_145728 to VT), the Swiss National Science Foundation (310030_143767 and 31003A_162609 to VT), the University of Basel and the Forschungsfonds of the University of Basel (RZ).

## MATERIALS AND METHODS

### Mouse strains and husbandry

The mouse models used in our experiments include: *Hes5::GFP BLBP::mCher* (Ables et al., 2010; Basak et al., 2012; Breunig et al., 2007; Ehm et al., 2010; Giachino et al., 2014; Imayoshi et al., 2010; Lugert et al., 2010; Lugert et al., 2012; Nyfeler et al., 2005) *Hes5::CreER^T2^, Notch2::CreER^T2-SAT^, Rosa26R-CAG::GFP, Rosa26R-CAG::tdTomato, Notch2^flox/flox^* and *Rosa26R-CAG::floxed-STOP-Notch2ICD* (Basak et al., 2012; Basak and Taylor, 2007; Besseyrias et al., 2007; Fre et al., 2011; Lugert et al., 2012; Schouwey et al., 2007). To confirm the expression of Notch2, we crossed *Notch2::CreER^T2-SAT^*, together with *Rosa26R-CAG::tdTomato*; to generate *Notch2* conditional knockout mice (*Notch2* CKO) and follow Hes5-expressing NSCs and their progeny, we crossed *Hes5::CreER^T2^* together with *Rosa26R-CAG::GFP* and *Notch2^flox/flox^*; to generate Notch2ICD constitutively expressed mice, we crossed *Hes5::CreER^T2^* together with *Rosa26R-CAG::GFP* and *Rosa26R-CAG::floxed-STOP-Notch2ICD*. Mice were genotyped by PCR of toe DNA. According to Swiss Federal and Swiss Veterinary office regulations, all mice were bred and kept in a specific pathogen-free animal facility with 12 hours day-night cycle and free access to food and water. All procedures were approved by the Basel Cantonal Veterinary Office under license numbers 2537 and 2538 (Ethics commission Basel-Stadt, Basel Switzerland).

### Tissue Preparation and Immunostaining

8-9 week old adult mice and 59-60 week old aged mice were injected daily intraperitoneal with 2 mg Tamoxifen in 100 μl sunflower oil for five consecutive days and sacrificed 1, 2, 21, 100 days or 10 months after the end of the Tamoxifen treatment. At different chase time points, mice were injected with ketamine/xylazine/acepromazine solution (130 mg, 26 and 4 mg per kg body weight, respectively) for deep anesthetization. The nesthetized mice were perfused with ice-cold 1x PBS, followed by PBS-buffered 4% paraformaldehyde (PFA). Brains were isolated, post-fixed overnight in PBS-buffered 4% PFA, and then immersed with 30% sucrose at 4°C overnight. The cryoprotected brains were frozen in OCT (TissueTEK) and cut as 30 μm floating coronal sections by cryostat (Leica).

For immunostaining, sections were blocked at room temperature for 30 minutes with 10% normal donkey serum (Jackson Immunoresearch) in PBS containing 0.4% TritonX-100. Afterwards, sections were incubated overnight at 4°C with the primary antibodies, which were diluted in PBS containing 2.5% normal donkey serum and 0.4% Triton X-100. Then sections were washed with PBS and incubated at room temperature for 2 hours with the corresponding secondary antibodies in 5% donkey serum blocking solution and stained with DAPI (1 μg/ml). After PBS wash, stained sections were mounted on glass slides (SuperFrost, Menzel), protected in DABCO anti-fading agent and visualized using a Zeiss ApoTome.2 microscope. For PCNA staining, the antigen was recovered at 80°C for 20 minutes in Sodium Citrate (10 mM, pH6.0) before primary antibody incubation.

### Stereotactic Injection of Virus Particles

*Adeno-gfap::Cre* and Retro pMI-NLSCre virus were produced respectively as previously described (Merkle et al., 2007; Zhao et al., 2006). Adult (8-10 weeks old) mice were anesthetized in a constant flow of isofluorane (3%) in oxygen and injected with Temgesic subcutaneously (0.05 mg/kg body weight) to relieve the pain. The head of the mice was fixed on a stereotaxic apparatus (David Kopf instruments), and the skull was exposed by an incision in the scalp and a small hole (Φ1mm) drilled through the skull. The *Adeno-gfap::Cre* virus or Retro pMI-NLSCre virus was injected in the dentate gyrus using a sharpened borosilicate glass capillaries at the stereotaxic coordinates anterior/posterior -2 mm; medial/lateral 1.5 mm relative to Bregma and -2 mm below the skull. Mice were sutured and monitored after surgery, and were then analyzed 21-days post-injection by immunostaining.

### Hippocampal Neural Stem Cell Isolation and FACS

*Hes5::CreER^T2^ Rosa26R-CAG::GFP* and *Hes5::CreER^T2^ Notch2^flox/flox^ Rosa26RCAG::GFP* 8-week old mice were injected with Tamoxifen for five consecutive days and sacrificed 1 day after the last injection. NSCs were isolated as previously described (Rolando et al., 2016). Briefly, brains were dissected in L15 Medium (GIBCO) and sectioned 300 μm thick using a McIllwains tissue chopper. The dentate gyrus was micro-dissected from the rest of the hippocampus under a binocular microscope avoiding contamination with tissue from the subventricular zone and molecular layer, digested with a papain based solution and mechanically dissociated. Cells were washed with L15 medium (Gibco, Invitrogen), filtered through a 40 μm cell sieve (Miltenyi Biotec) and sorted by forward and side-scatter for live cells (control) and gated for GFP-negative (wild type levels) or GFP-positive populations with a FACSaria III (BD Biosciences). Live GFP-positive cells were used for gene expression analysis. Each genotype contains 3 independent samples, each sample are pooled cells from 4-5 mice.

### RNA-seq and Data Analysis

Total RNA was isolated from sorted DG NSCs using Trizol reagent (Life Technologies). Library preparation and transcriptome sequencing were performed by sequencing facility at D-BSSE (Basel), according to the Smart-seq2 protocol (Invitrogen). Sequencing data was analyzed by using an unbiased Suvrel Method based on supervised relevance learning in order to obtain the lists of differentially regulated genes (Boareto et al., 2015). By defining a specific cost function, we selected the genes based on how much they increase the distance between samples of different experimental conditions and approximate samples of same experimental conditions. Reads were mapped to genome using Bowtie2 (Langmead and Salzberg, 2012) and transcript identification and quantification were performed using HTSeq algorithm (Anders et al., 2015). Source code is available at: https://git.bsse.ethz.ch/marcelob/Notch2KO. The RNA-seq data are available at GEO under the accession number: GSE116773.

### Hippocampal Adult NSC Cultures

DG NSCs were isolated from 8-week-old mice as described in the part of *Hippocampal Neural Stem Cell Isolation and FACS*. Cells were plated in dishes (Costar) coated with 100 µg/ml Poly-L-Lysine (Sigma) and 1 µg/ml Laminin (Sigma) in neural progenitor culture medium containing DMEM: F12 (Gibco, Invitrogen), 2% B27 (Gibco, Invitrogen), FGF2 20 ng/ml (R&D Systems), EGF 20 ng/ml (R&D Systems).

### Myc-tagged Notch2ICD ChIP

*Rosa26R-CAG::GFP* control and Notch2ICD overexpressing NSC cultures were harvested 60 hours after Cre infection. 1 ml 10x fixation buffer (0.7 ml dilution buffer (1 M NaCl, 50 mM Tris, pH8, 10 mM EDTA, 5 mM EGTA) and 0.3 ml 16% Formaldehyde (ThermoFisher, Methanol-free, 28906)) were added per each 10 cm dish (about 5 million cells), and cells were kept for 10 min at room temperature. 1.1 ml 2.5 M glycine was added and cells were incubated for another 10 min. Supernatant was discarded and cells were washed with 10 ml ice cold PBS. Per each plate, cells were scraped with 2 ml ice cold PBS, transferred to a 15 ml Falcon tube, and centrifuged at 500 g for 5 min at 4°C. Cell pellets were washed in 15 ml Solution A (10 mM Tris pH8, 0.25% Triton, 10 mM EDTA, 0.5 mM EGTA), kept on ice for 5 min, and centrifuged 500g for 5 min at 4°C. The pellets were further washed in 15 ml Solution B (10 mM Tris pH8, 200 mM NaCl, 1 mM EDTA, 0.5 mM EGTA) twice, and centrifuged at 500 g for 5 min at 4°C. The pellets were dissolved in 1 ml Sonication buffer (10 mM Tris pH8, 1 mM EDTA, 0.5 mM EGTA) and sonicated with Covaris at 140 w/5% duty factor/200 cycles per burst for 15 min. The chromatin solution was transferred to 1.5 ml tubes and centrifuged for 15 min at the maximum speed. The supernatant was then transferred to new tubes (500 μl aliquots), and adjusted to RIPA conditions by addition of 100 μl 10% TritonX100, 100 μl 1% Na-deoxycholate, 100 μl 1% SDS, 100 μl 1.4 M NaCl, and 100 μl 50 mM Tris pH8. Protease inhibitors (Roche complete; or 2 μg/μl Pepstatin, Leupeptin, Aprotinin, 1 mM PMSF) were added. Samples were centrifuged for 10 min with 13000 rpm at 4°C. The supernatants were transferred to new siliconized tubes. 100 μl of each chromatin batch was used for input control (10% input). 3 μg anti-Myc tag (Abcam, goat, ab9132, ChIP-grade) antibodies were added and tubes were rotated over night at 4°C. Protein G Dynabeads were equilibrated with RIPA buffer. ChIP samples were centrifuged for 5 min at high speed, transferred to properly labeled tubes containing 40 μl equilibrated Dynabeads, and rotated for 2-4 hours at 4°C. The Dynabeads/immune complexes were washed with RIPA buffer (10 mM Tris pH8, 1 mM EDTA, 140 mM NaCl, 1% Triton, 0.1% SDS, 0.1% Na-deoxycholate, Roche protease inhibitors) 5 times, with LiCl buffer (10 mM Tris, pH8, 250 mM LiCl, 1 mM EDTA, 0.5% NP40, 0.5% Na-deoxycholate) once and with TE buffer (10 mM Tris, pH8, 1 mM EDTA) twice. After the last washing step, 100 μl freshly prepared Elution buffer (1% SDS, 0.1 M NaHCO3) was used to re-suspend Dynabeads by pipetting up and down. The beads were incubate at 65°C for 15 min with gentle agitation (800 rpm) and centrifuged briefly, and the supernatant was transferred to a new tube. Elution was repeated once and elutes were combined. 12 μl of 5 M NaCl was added into 200 μl eluate. Reverse crosslinking was performed by incubation at 65°C overnight (>=6hrs). DNA was then treated by RNase and Proteinase K, purified by phenol/chloroform extraction and ethanol precipitation, and prepared for qPCR analysis.

### Gene overexpression and knockdown by Nucleofection

*Rosa26R-CAG::GFP* control and *Rosa26R-CAG::floxed-STOP-Notch2ICD* adult DG NSC cultures were nucleofected according to the mouse neural stem cell kit instructions (Lonza). Briefly, DG NSCs were dissociated with trypsin and resuspended in the nucleofector solution to the final concentration of 4×10^6^ cells/100 μl. For Id4 overexpression, DG NSCs suspensions were combined with either pLVX-IRES-tRFP or pLVX-Id4-IRES-tRFP plasmids. For Id4 knockdown, DG NSCs suspensions were combined with either 200 pmol antisense LNAGapmeRs against Id4 or negative control LNAGapmeRs (Exiqon). A pCAG::mCherry expression vector was added to trace transfected NSCs. Then DG NSC suspensions were nucleofected with Lonza Nucleofector 4D device (program DN-100) according to the manufacturer’s instructions. Transfected NSCs were immediately transferred into NSC culture medium and plated on poly-L-Lysine/Laminin coated dishes or coverslips. For the Id4 overexpression, cells were fixed 48 hours later for gene expression and proliferation analysis. For Id4 knockdown, 24 hours after nucleofection, DG NSCs were infected with *Adeno-Cre* virus and were fixed 48 hours later for proliferation analysis. To analyze RNA levels of *Id4*, GFP mCherry double positive cells were sorted 24 hours after *Adeno-Cre* virus infection for RNA preparation and RT-qPCR.

### RT-qPCR and ChIP-PCR

For RT-qPCR test, total RNA was isolated from sorted DG NSCs or NSC cultures by using Trizol reagent (Life Technologies) according to the manufacturer’s instructions. Low amount RNA isolated from sorted DG NSCs was processed and amplified by using REPLIg^®^ WTA Single Cell Kit (Qiagen). High amount RNA isolated from NSC cultures was reverse-transcribed to cDNA by using SuperScript™ IV VILO™ Master Mix with ezDNase™ Enzyme Kit. qPCR reactions were performed on a Rotor-Gene Q PCR machine (Qiagen) by using PowerUp™ SYBR^®^ Green Master Mix (Life Technologies).

For ChIP-PCR, primers were designed according to Rbpj binding motif (GTGGGAA). Input DNA and pull-down DNA by myc antibody were amplified and quantified by qPCR. The final fold enrichment of Notch2ICD binding sites were calculated by the binding enrichment (pull-down/Input) in the Notch2ICD samples relative to the *Rosa26R-CAG::GFP* control samples. Primers for RT-qPCR and ChIP-PCR reactions are described in Table S7.

### BrdU and EdU Administration

For *in vivo* labeling, BrdU was dissolved in the drinking water at 1 mg/ml with 1% sucrose. Aged mice (60-week-old) were fed with BrdU-water from the beginning of the Tamoxifen treatment to one week after the treatment (12 days in total). Mice were sacrificed 21 days after the end of the Tamoxifen treatment and perfused transcardially with 4% PFA in PBS. Cryo-protected tissue was sectioned at 30 μm and immunostained as floating sections by incubation with BrdU antibodies.

For *in vitro* labeling, half of the NSC media was replaced with fresh media containing 20 μM of BrdU or EdU to obtain a final concentration of 10 μM. Cells were incubated for 4 hours before fixation. Next, media was removed and 1 mL of 4% PFA in PBS was added to each well containing the coverslips. Cells were incubated for 15 minutes at room temperature. Fixation buffer was removed and the cells in each well were washed twice with 1 mL of 3% BSA in PBS. Wash solution was removed and 1 mL of 0.5% Triton^®^ X-100 in PBS was added to each well and incubated at room temperature for 20 minutes.

DNA hydrolysis was performed before BrdU staining. Cells or tissue sections were fixed with 2N HCl at 37°C for 15 min, and then neutralized with 0.1 M sodium borate buffer pH 8.5 for 30 minutes at room temperature. Samples were wash three times in PBS and continued with immunostaining for BrdU antibody. EdU detection was performed according to the instruction of Click-iT EdU Alexa Fluor 647 Imaging Kit.

### Quantification and statistical analysis

Images of immunostainings were captured and processed with or without Z-stacks on a Zeiss Apotome2 microscope. Stained cells were counted manually using ImageJ (Schindelin et al., 2012). Data are presented as averages of indicated number of samples. The average number of each sample was determined by minimum of six DG sections per animal or 10 views per cell culture staining. Data representation and statistical analysis were performed using GraphPad Prism software. Two-tailed unpaired Student’s t test was used to compare two samples. Percentages were transformed into their arcsin values to perform statistical analysis. The number of replicates (n) is indicated in the corresponding quantification tables in the supplemental items. Statistical significance is determined by p values (*p <0.05, **p <0.01, ***p < 0.001) and error bars are presented as SEM.

